# Predicting Parallelism and Quantifying Divergence in Experimental Evolution

**DOI:** 10.1101/2020.05.13.070953

**Authors:** William R. Shoemaker, Jay T. Lennon

## Abstract

The degree that the environment determines what genes contribute towards adaptation is a fundamental question in microbial evolution. Microbial populations are often experimentally passaged in different environments and sequenced in order to identify candidates for adaptation in a particular environment. However, there remains the need to develop an appropriate statistical framework to identify genes that acquired more mutations in one environment over the other (i.e., divergent evolution). Here we demonstrate how the evolutionary outcomes among replicate populations in the same environment, known as parallel evolution, can be leveraged to construct an intuitive statistical test for identifying the genes that contribute towards divergent evolution. To accomplish this task, we examined publicly available evolve-and-resequence experiment datasets and found that the distribution of mutation counts among genes can be predicted using an ensemble of independent Poisson random variables. Building on this result, we propose that the degree of divergent evolution at a given gene between populations from two different environments can be modeled as the difference between two Poisson random variables, known as the Skellam distribution. We then propose and apply a statistical test to identify specific genes that contribute towards divergent evolution. IMPORTANCE: There is currently no existing framework that can be leveraged to identify genes that contribute towards divergent evolution in microbial evolution experiments. To correct for this absence, we investigated the distribution of mutation counts among genes in order to identify an appropriate null model. Our observations suggest that divergent evolution within a given gene can be modeled as the difference in the total number of mutations observed between two environments. This quantity is described by a probability distribution known as the Skellam distribution, providing an appropriate statistical test for researchers seeking to identify the set of genes that contribute towards divergent evolution in evolution experiments.

## Observation

Biologists have long been fascinated by the degree to which evolution is repeatable^1^. Independently evolving populations frequently evolve similar genotypes and phenotypes, a phenomenon known as parallel evolution^2,3^. Parallel evolution is particularly prevalent among microorganisms. The rise of evolve-and-resequence experiments as high-throughput screens for adaptation^4^ has allowed researchers to identify recurrent mutations across replicate populations^4,5^, paring down the vast number of potentially adaptive mutations into those that putatively confer the largest fitness benefits. Furthermore, evolve-and-resequence experiments have revealed that the outcomes of evolution are often conditional on the ancestral genotype of a microbial population^6–10^ or the environment in which it was maintained^11–15^, a phenomenon known as divergent evolution.

The ease in which evolve-and-resequence experiments can be performed comes with the drawback that there is comparatively little statistical direction on how the contributors of divergent evolution should be identified. In recent years, models that coarse-grain over molecular details have been remarkably successful in identifying general microbiological principles^16^. These models, and the underlying motivation to develop straightforward interpretations of biological phenomena, raises the question of whether there is an intuitive way in which contributors towards divergent evolution can be identified. To address this issue, we first determined the extent that we can predict patterns of parallel evolution at the gene level using a straightforward statistical model and publicly available data. Building on these results, we formulated and tested an interpretable model of divergent evolution at the gene level.

### Predicting genetic parallelism among replicate populations

The task of identifying genes that contribute towards divergent evolution can be viewed as the equivalent of identifying genes that undergo a greater degree of parallel evolution in one environment relative to another (Fig. 1). This observation suggests that it is necessary to first identify an appropriate model of parallel evolution within a single environment in order to develop a model of divergence. Given that the per-generation probability of acquiring a mutation at a given gene is low and the number of generations is large, it is reasonable to assume that a given gene acquires mutations as a Poisson process. We can model the sampling distribution of this process as the probability of observing *n*_*i, j*_ mutations in the *i*th gene within a population that acquired a total of *n*_tot, *j*_ mutations as

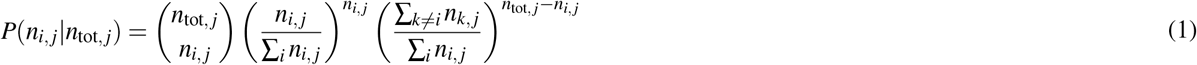

**Figure 1.**
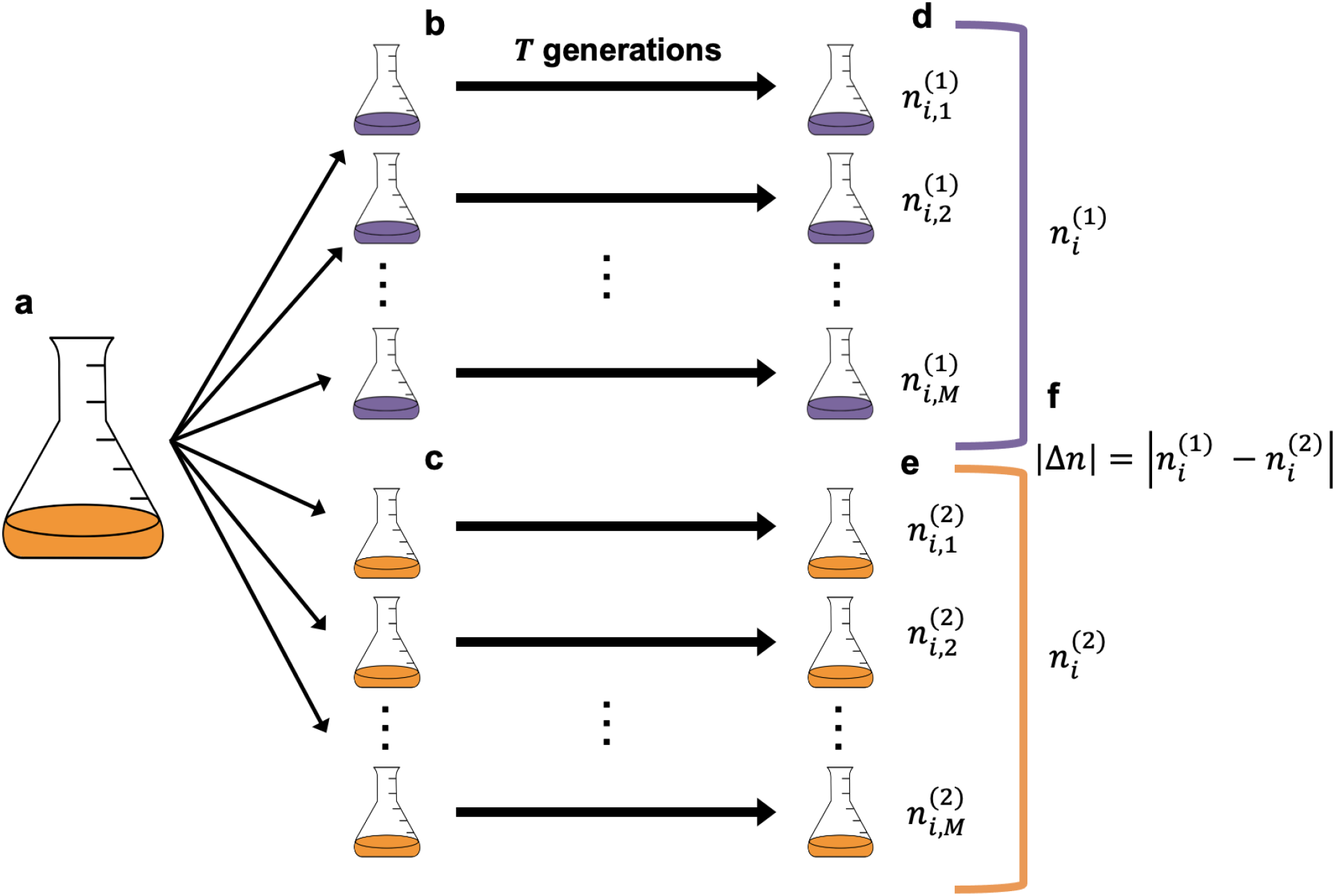
**a)** A typical evolve-and-resequence experiment is performed by splitting a culture grown from a single colony inoculate into replicate flasks constituting one or more environment (e.g., purple or orange) and propagating the culture over time. **b**,**c)** After a given number of generations has elapsed, replicate populations are often sequenced, allowing for the number of *de novo* mutations at a given gene to be calculated. **d**,**e)** The degree of parallel evolution within each environments is quantified by taking the sum of mutation counts across replicate populations for a given gene, **f)** while the degree of divergent evolution is quantified by taking the absolute difference in mutation counts between environments (|Δ*n*|)

We can then determine whether we can predict statistical patterns from empirical data using Eq.1. Given that mutation count data from evolve-and-resequence experiments are often sparse, it is natural to calculate the proportion of populations that have at least one mutation in a given gene (i.e., *occupancy, o*_*i*_^17^) and compare our empirical estimate to an expected value by averaging over *M* replicate populations

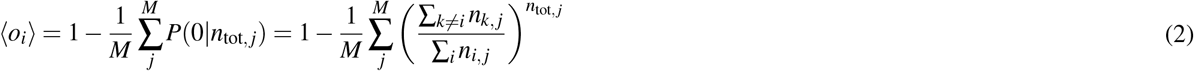

To test our prediction, we calculated ⟨*o*_*i*_⟩ from Eq. 2 on nonsynonymous mutation count data from an evolve-and-resequence experiment with 115 replicate populations of *E. coli*^11^. We found that our model does a reasonable job capturing the observed occupancy (Fig. 2a) with a mean absolute error (MAE) of ∼ 0.008. However, while MAE decreased with an increasing number of replicate populations, it ultimately saturated (Fig. 2b). The fact that it does not reach zero suggests that features not incorporated into our model such as non-independence among genes may be necessary to fully explain the distribution of mutation counts.

**Figure 2.**
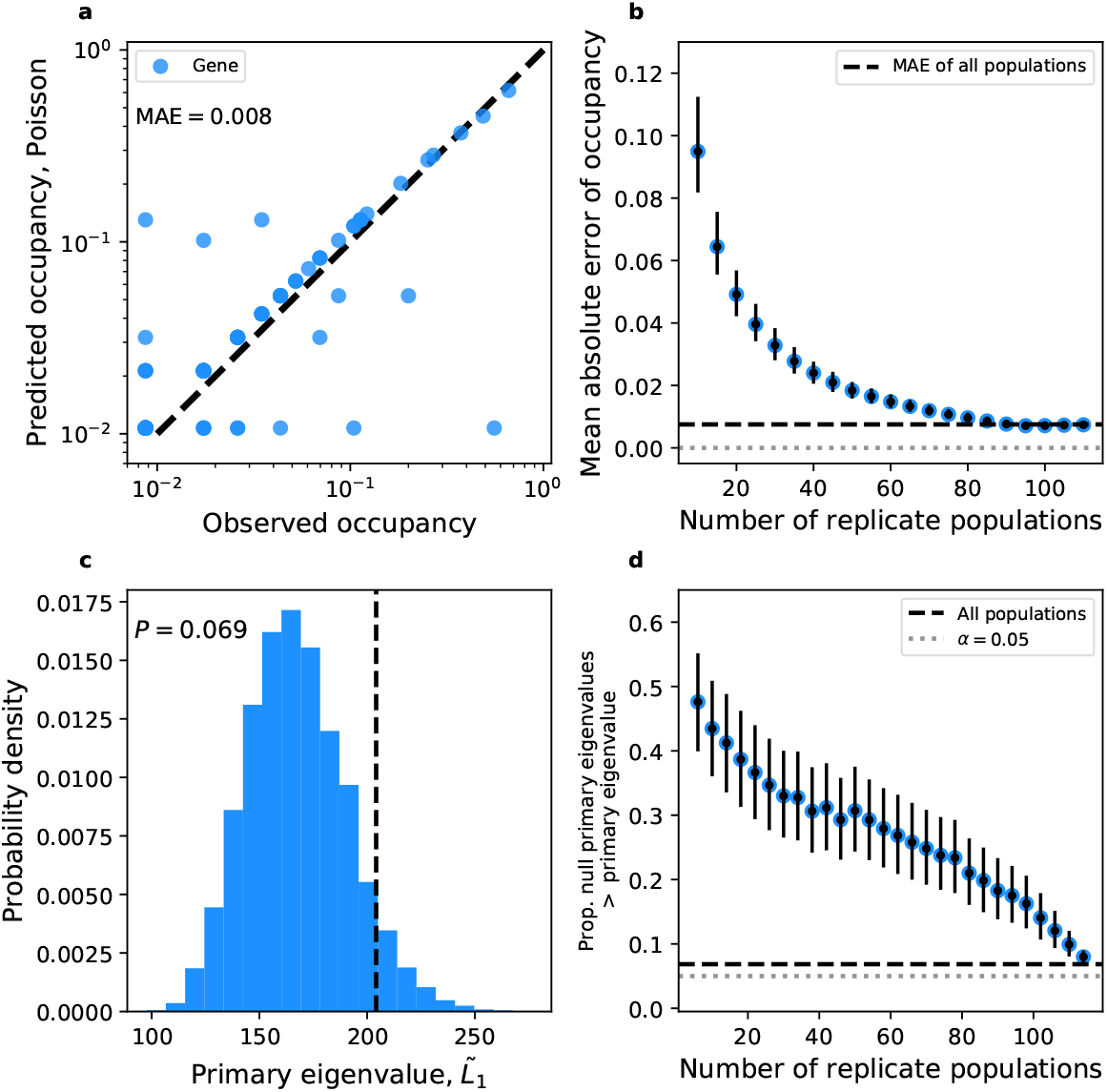
**a)** Using the Poisson distribution, we can predict the occupancy of nonsynonymous mutations for a given gene among 115 replicate *E. coli* populations. **b)** The amount of error rapidly decreases as the number of replicate populations increases. **c)** The degree of covariance in a gene-by-population matrix can be summarized by the primary eigenvalue (dashed black line). By generating null count matrices, we can calculate a null distribution of primary eigenvalues and calculate a *P*-value. **d)** By subsampling replicate populations without replacement, we can calculate the fraction of observed primary eigenvalues greater than the null.

To determine whether non-independence among genes was necessary to incorporate in our model, we tested whether we could detect signals of covariance in our data. Because the number of genes that acquired mutations in an experiment can number in the hundreds and mutation count data is sparse, attempting to estimate individual covariances would be unreasonable. Instead, we can estimate a global signature of covariance and compare it to a null distribution (Methods). While the global signal of covariance increased with the number of replicate populations, it was weak for values typical of most evolution experiments (5-20 populations; Fig. 2a,b) and was only borderline significant when all 115 replicate populations were included (*P* = 0.072).

### Identifying contributors of divergent evolution between a pair of environments

The success of the multivariate Poisson in describing the distribution of mutation counts within a given environment and the overall weak signals of covariance provide the justification necessary to model the distribution of mutation counts among genes as an assemblage of effectively independent variables. We can then model divergent evolution at a given gene as the difference in two independent Poisson rates. In terms of mutation counts, we can identify the meaningful variable as the absolute difference in mutation counts between two environments for a given gene (|Δ*n*|). The distribution of |Δ*n*| has been previously derived and is known as the Skellam distribution^18^. Starting with the null Poisson rates for each treatment 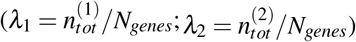, we define the probability mass function of the absolute value of 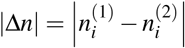 as

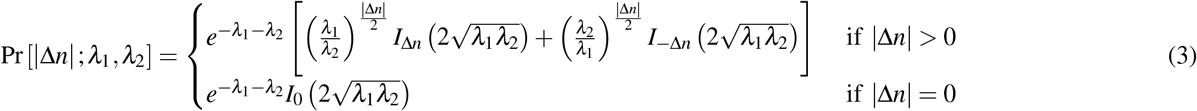

where *I*_Δ*n*_(·) is a modified Bessel function of the first kind. Building on a previous approach developed to identify contributors of parallel evolution^19^, we can define the *P*-value as

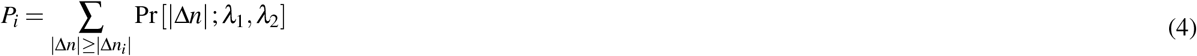

To reduce the number of tests we can only calculate *P*-values for |Δ*n*| ≥ *n*_*min*_, where the expected number of genes with |Δ*n*| ≥ *n*_*min*_ and *P*_*i*_ ≤ *P* is

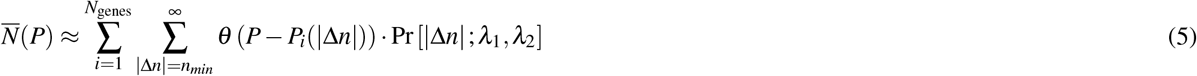

where *θ* (·) is the Heaviside step function. We can then compare this number to the observed number of genes *N*(*P*), defining a critical *P*-value (*P*^∗^) for a given FDR *α* as

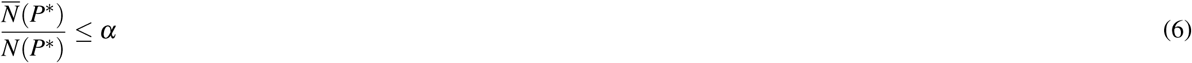

We then applied this approach to an experiment with replicate populations for four treatments^12^. We were able to identify genes that were consistently enriched for nonsynonymous mutations within a given treatment across all pairwise treatment comparisons (Table S1), largely agreeing with the conclusions of the original study^12^.

## Concluding Remarks

In this study, we investigated the distribution of mutation counts in evolve-and-resequence experiments. We found that a Poisson model sufficiently explains the distribution of mutation counts across genes. Using this result, we proposed that the difference in Poisson rates between treatments (i.e., the Skellam distribution) can be used to identify genes that contribute towards divergent evolution. These results can serve as a useful tool for analyzing the results of evolve-and-resequence experiments.

## Methods

### Predicting and quantifying parallelism

To determine the degree that we can predict statistical patterns of parallel evolution, we used a publicly available dataset of one of the largest microbial evolve-and-resequence experiments. In this experiment, 115 replicate populations of *Escherichia coli* were serially transferred for 2,000 generations at 42.2 °C^11^. A single colony was isolated from each replicate population and sequenced.

To test for a global signal of covariance between genes, we merged all nonsynonymous mutations from all replicate populations into a population-by-gene count matrix. To account for gene size as a covariate, we corrected the number of mutations in all empirical data by calculating the excess number of mutations (i.e., *multiplicity*) 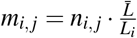, where *L*_*i*_ is the number of nonsynonymous sites in the *i*th gene and 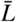 is the mean of all genes in the genome^19^. To determine whether covariance can be reliably detected a given level of replication we estimated the largest normalized eigenvalue^20,21^, defined as

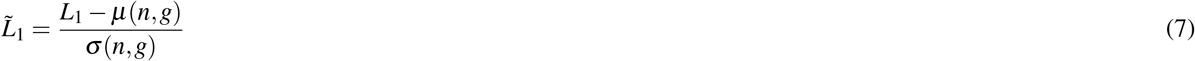

where *L*_1_ is normalized as 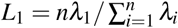 to sum to *n* and

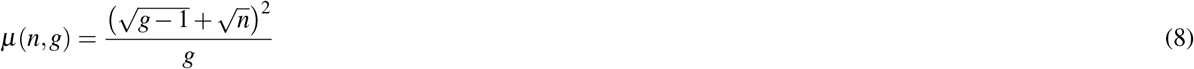

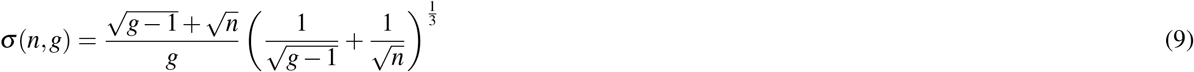

As *n, g* → ∞ and 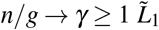 tends towards a Tracy-Widom distribution^21,22^. Though these limits can be relaxed^20,23^. A null distribution of 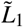 was obtained by randomizing combinations of mutation counts constrained on the total number of mutations acquired within each gene across treatments and the number of mutations acquired within each treatment. Randomization was performed using a Python implementation^24^ of the ASA159 algorithm^25^.

## Available Code and Data

Instructions to obtain public data and code to reproduce our analyses are on GitHub: https://github.com/LennonLab/ParEvol.

## Acknowledgments

This work was supported by US Army Research Office Grant W911NF-14-1-0411.

## Supplemental material

**Supplementary Table S1.** Using Eq. 4, we calculated a *P*-value of divergent evolution for each pairwise treatment comparison for each gene. To identify candidates of adaptation that are unique to a given treatment, we identified the set of genes that were significantly enriched for nonsynonymous mutations within a given treatment for all pairwise treatment comparisons.

| Treatment | Locus tag | Function |
| --- | --- | --- |
| High carbon, bead | BCEN2424_RS08045 | Type 1 fimbrial protein |
| High carbon, planktonic | BCEN2424_RS08125 | Lysine N(6)-hydroxylase |
| Low carbon, bead | BCEN2424_RS16880 | MFS transporter |
|  | BCEN2424_RS25830 | FkbM family methyltransferase |
| Low carbon, planktonic | BCEN2424_RS03065 | IclR family transcriptional regulator |
|  | BCEN2424_RS04940 | YicC family protein |

